# Automated Segmentation and Tractography of Brainstem and Subcortical White Matter

**DOI:** 10.64898/2026.06.09.731210

**Authors:** Kurt G Schilling, Gaurav Rudravaram, Antoine Theberge, Matthew Amandola, Michael E. Kim, Kathryn L. Humphreys, Laurie Cutting, Derek Archer, Timothy J. Hohman, Angela L. Jefferson, Lori L. Beason Held, Murat Bilgel, Alzheimer’s Disease Neuroimaging Initiative, The BIOCARD Study Team, Maxime Chamberland, Maxime Descoteaux, Laurent Petit, Francois Rheault, Bennett Landman

## Abstract

Diffusion MRI enables noninvasive mapping of human white matter pathways, but automated segmentation methods have largely focused on large association, projection, and commissural bundles. Brainstem and subcortical pathways supporting basal ganglia, cerebellar, limbic/reward, sensory, and homeostatic functions remain underrepresented in large-scale connectomic analyses. To address this gap, we adapted BundleParc, a recently introduced bundle-parcellation architecture, into an automated pipeline for direct segmentation and along-tract parcellation of 97 subcortical and brainstem white matter pathways. The model was trained on a curated reference dataset derived from Human Connectome Project diffusion MRI using anatomy-guided tractography, explicit inclusion and exclusion criteria, automated outlier filtering, and manual quality assurance. Operating directly on native-space fiber orientation distributions, the algorithm successfully recovers these intricate anatomical trajectories and ordered parcellations. We show the model generalizes to diverse external datasets spanning development, aging, and clinical cohorts, with performance varying across cohorts and acquisition protocols. The released container, trained model, population atlas, reference streamlines, and quality assurance outputs provide a resource for quantitative studies of brainstem and subcortical pathways.

## 1. Introduction

Diffusion MRI (dMRI) fiber tractography has transformed our understanding of the human connectome, facilitating the non-invasive, in vivo mapping of complex neural architectures [1]. dMRI measures the directional dependence of water diffusion in tissue, allowing local fiber orientations to be estimated and followed from voxel to voxel to reconstruct white matter pathways. Over the past two decades, significant progress has been made in the automated segmentation (i.e., virtual dissection) and quantitative analysis of white matter pathways[2–7]. However, current analytical frameworks have predominantly focused on the large, long-range association, projection, and commissural bundles of the cerebral mantle. In comparison, the dense, compact, and anatomically complex white matter of the brainstem and subcortex [8, 9] has received much less attention and remains relatively underexplored[10].

The pathways within the brainstem and subcortex include basal ganglia loops, cerebellar outflow systems, limbic and reward connections, and the deep intrinsic tracts that support autonomic and homeostatic regulation. Together, these white matter systems contribute to voluntary motor control, somatosensory processing, nociception, arousal, and other functions that are central to both neurologic and psychiatric disease [9]. Many of these pathways also run through regions that are common targets for deep brain stimulation and other stereotactic interventions [11]. Yet, their study in vivo is limited by a combination of anatomical and imaging constraints [12]. Functionally distinct systems are interleaved within a small volume, and many relevant pathway origins, terminations, and trajectories lie in small or low-contrast nuclei and subregions. The pathways themselves are small, closely packed, and frequently cross, merge, or fan in regions where conventional diffusion MRI is limited by spatial resolution, tissue contrast, distortion, and physiological motion [13]. Overall, this makes these pathways difficult to define and segment reproducibily in MRI workflows.

Optimistically, a growing body of work has shown that many of these pathways can be delineated with tractography using high-quality data and anatomy-informed expert-guided protocols [11, 14, 15]. Connectome-based and probabilistic atlases have mapped brainstem trajectories in vivo [16–19], tractography pipelines have reconstructed selected brainstem tracts [20–25], and recent work has begun to automate segmentation of both brainstem nuclei and sets of brainstem white matter bundles[26, 27]. At the same time, high-resolution subcortical connectomics is rapidly expanding the anatomical reference base (i.e., the start and end points) for these connections[28]. A key gap, however, is the lack of a broad, subject-specific, automated framework that can jointly segment deep pathways across multiple subcortical and brainstem systems and do so in a way that is practical for *in vivo* diffusion MRI datasets.

To address this gap we constructed an end-to-end automated pipeline to segment 97 white matter pathways across multiple subcortical and brainstem systems. Using curated anatomical-tractographic reference labels derived from Human Connectome Project (HCP) [29] diffusion MRI, we trained a bundle-parcellation model to reproduce these pathways as subject-specific bundle masks and within-bundle parcellations directly from fiber orientation distributions [30].

We show that the framework not only maps these pathways accurately but also generalizes across heterogeneous external datasets that vary in age, disease context, acquisition scheme, and image quality. The broader aim is to make these comparatively undercharacterized pathways accessible for quantitative studies of development, aging, disease, and neuromodulation-relevant anatomy.

## 2. Methods

### 2.1. Overview of Study Design

We developed an automated framework for segmentation and parcellation of 97 brainstem and subcortical white matter pathways (full pathway names and abbreviations are given in **Appendix A**). The workflow comprised four stages. First, we defined and curated the pathway set in HCP diffusion MRI using semi-automated tractography with manual quality assurance, yielding silver-standard reference labels in the form of bundle masks and ordered within-bundle parcellations. Second, these reference labels were used to train a recently designed BundleParc-based segmentation network[30] that predicts subject-specific pathways from fiber orientation distributions. Third, we quantified performance on held-out subjects using both binary segmentation and parcellation metrics. Fourth, we assessed generalizability across external datasets spanning broad variation in age, disease context, acquisition scheme, and image quality. When streamline representations were desired for visualization or downstream quantitative analysis, we additionally performed tractography constrained by the predicted segmentations and endpoint parcels.

### 2.2. Imaging Datasets

#### 2.2.1. HCP training/validation/test data

Reference labels for model development were derived from the HCP Young Adult dataset [29]. We began with 1050 subjects (age range 22-35) with structural and diffusion MRI. The diffusion acquisition provided 1.25 mm isotropic data with three shells at b = 1000, 2000, and 3000 s/mm² (90 diffusion directions per shell), which enabled detailed tract reconstruction and anatomical curation. Diffusion data were preprocessed with the HCP minimal preprocessing pipeline [31], which included susceptibility distortion, motion, and eddy current corrections and registration to the subject’s AC-PC-aligned structural space. Multi-shell multi-tissue constrained spherical deconvolution was used to extract the fODF (MRtrix3 [32]; dwi2response dhollander [33, 34]; dwi2fod msmt [35]). This dataset served as the basis for semi-automated tractography, manual quality assurance, and the silver-standard reference labels used for training and evaluation.

#### 2.2.2. External datasets for generalizability

Generalizability was assessed on 18 external diffusion MRI datasets acquired across multiple sites and spanning a broad range of ages, clinical populations, and acquisition protocols (datasets, acronyms, and citations given in **Appendix B**). These datasets included infant, developmental, young adult, and aging cohorts, as well as cohorts involving attention-deficit/hyperactivity disorder (ADHD), dyslexia, multiple sclerosis (MS), and mild cognitive impairment or Alzheimer disease (AD). Ten subjects were randomly selected from each dataset. Across cohorts, acquisitions varied in spatial resolution, isotropic versus anisotropic sampling, b value, number of diffusion directions, and single-shell versus multi-shell design. All external diffusion MRI data were preprocessed with PreQual (v1.0.8) [36] to correct susceptibility-induced echo-planar imaging distortion, head motion, and eddy current artifacts before model inference.

### 2.3. Pathway set and anatomical definitions

#### 2.3.1. Selection and reconstruction of 97 pathways

Pathway definitions were guided by prior anatomical and tractographic studies, primarily driven by work on brainstem tractography atlases [17, 27], subcortical deep brain stimulation circuitry [11, 37], functional circuitry [38], cerebellothalamic systems [22, 24], and tractography demonstrations of specific pathways [25]. Pathway definitions were developed with guidance from an expert neuroanatomist (L.P.).

Anatomical guidance was derived from several cortical and subcortical atlases, including FreeSurfer-derived[39] Desikan–Killiany [40] and Brodmann cortical labels, the CIT168 [41] and HybraPD [42] subcortical nuclei atlases, DISTAL, the DBS tractography atlas [11], Multi-contrast PD25 atlas [43], Thalamus Optimized Multi Atlas Segmentation (THOMAS) atlas [44], and the Zona Incerta atlas [45], together with manually defined regions of interest when an appropriate atlas label was not available. Atlas-derived regions of interest were transformed into each subject’s diffusion space using ANTs registration of the corresponding FA images, including rigid, affine, and nonlinear SyN stages (note, atlas registration is not required for inference on new or external datasets). Silver-standard bundle masks and parcellations were then generated from subject-specific tractography.

For each pathway, we defined anatomical rules specifying where streamlines were required to start, end, and pass, or do not pass through [46] - implemented with combinations of seed, stop, inclusion, and exclusion regions of interest. When appropriate, we evaluated both directional formulations of a pathway by reversing seed and stop regions, and also tested whole-brain seeding with paired endpoint regions enforced as inclusion constraints.

All candidate pathways were first filtered with automated outlier rejection using *scil_bundle_reject_outliers* in Scilpy[47]. We then performed exhaustive visual quality assurance across the full HCP Young Adult cohort and the full pathway set. For each subject-pathway pair, standardized multi-view QA screenshots were generated, together with the contralateral homolog. Reconstructions were reviewed for predefined failure modes, including anatomically implausible trajectories, incorrect origin or termination, false-positive collateral streamlines, missing or fragmented segments, marked left-right asymmetry, complete tracking failure, or persistent outlier streamlines. Each subject-pathway pair was assigned a pass or fail quality label. When false-positive connections or other clear anatomical errors were present, streamlines were inspected and edited manually in MI-Brain [48], and the reconstruction was reevaluated. A conservative quality assurance strategy was used throughout. Questionable bundles, clear failures, bundles with no streamlines, and persistent outliers were excluded from the final reference set. Only subject-pathway pairs that passed this quality assurance process were converted into reference masks and along-tract parcellation labels for model training, validation, and testing. Final pathway counts after quality assurance are provided in **Appendix C. Figure 1** shows representative reference pathways from a single subject, grouped by system for visualization.

**Figure 1.**
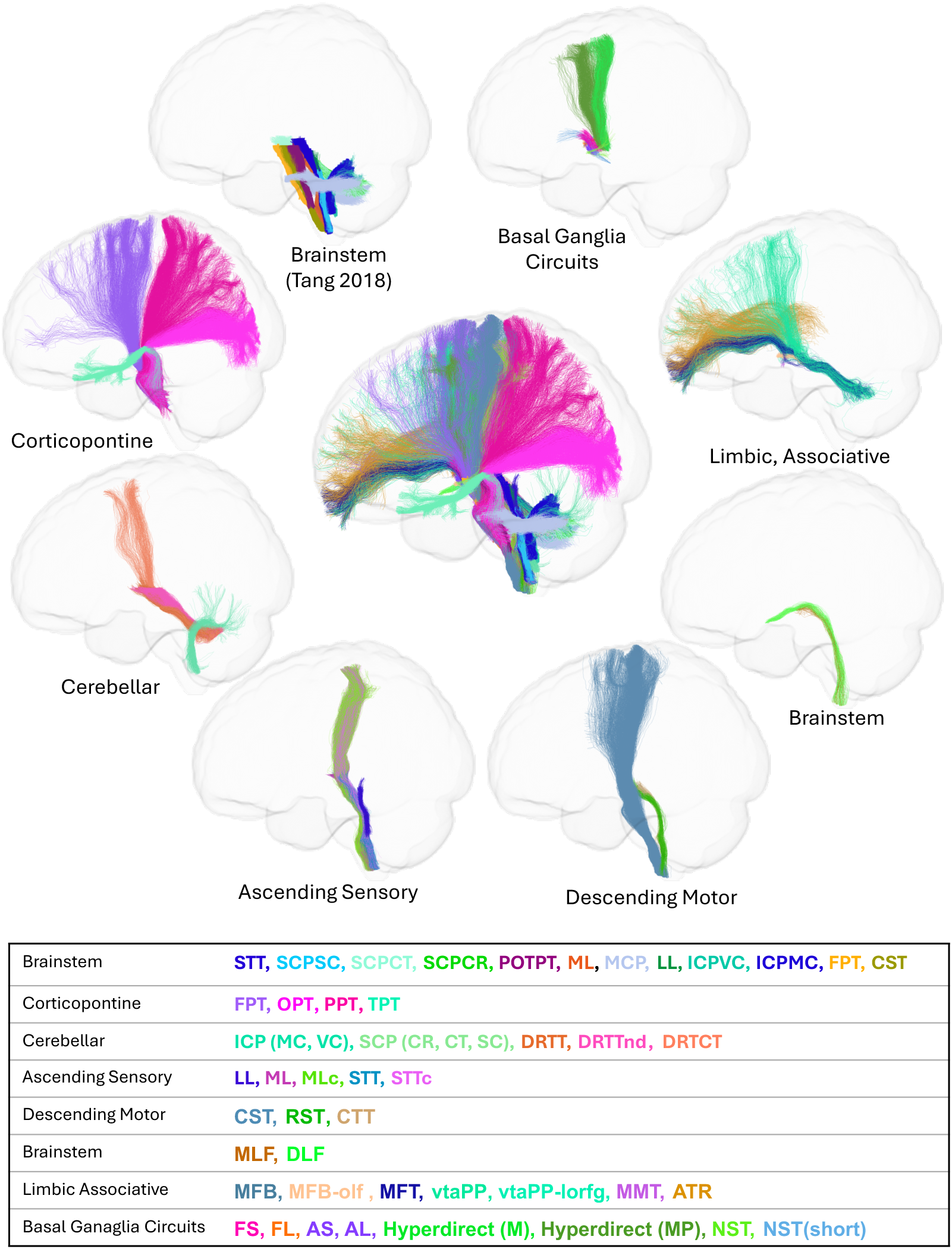
Representative reference pathways from a single HCP subject. Composite rendering of the curated pathway set used for silver-standard reference label generation. For bilateral pathways, one hemispheric example is shown to reduce visual clutter. Pathways are roughly grouped by location or functional systems for display and visualization.

#### 2.3.2. Multi-axial classification framework

There are several ways to group, organize, and interpret these pathways. We classified each pathway along four complementary axes - gross morphology, topographic location, physiological role, and participation in named circuits. This framework was used to organize the pathway set, to support presentation of the results, and future use by other groups.

The first axis, morphological class, describes the gross connectivity pattern of each pathway. In the present pathway set, pathways were classified as either projection or intrinsic. Projection pathways link cortex, deep nuclei, brainstem, cerebellum, or spinal cord across major compartments of the nervous system, whereas intrinsic pathways remain largely confined to the brainstem or to tightly coupled local brainstem circuits. The second axis, topographical subclass, describes whether a tract is superficial, deep, or peduncular according to its position relative to the ventrolateral brainstem surface, the tegmental core, and the cerebellar peduncles. The third axis, functional subclass, groups pathways by their dominant physiological role, including motor, sensory, cerebellar modulatory, autonomic or homeostatic, and limbic or reward functions. The fourth axis, network subclass, assigns each tract to a named circuit when appropriate, including pallidothalamic, hyperdirect, cerebello-thalamo-cortical, Papez, Guillain-Mollaret, and central homeostatic systems.

Pathway origins, terminations, classifications (and additional information) are presented in Tables 1-4, organized primarily around shared functional and network relationships (they do not correspond one-to-one with a single axis).

Table 1 contains the basal ganglia, limbic, and reward pathways that course through the pallidal, subthalamic, thalamic, nigral, and ventral tegmental regions. This group includes pallidothalamic and pallidosubthalamic projections, cortico-subthalamic hyperdirect pathways, ventral tegmental projections, and limbic or diencephalic connections that link basal ganglia output with forebrain and reward-related systems.

**Table 1.**
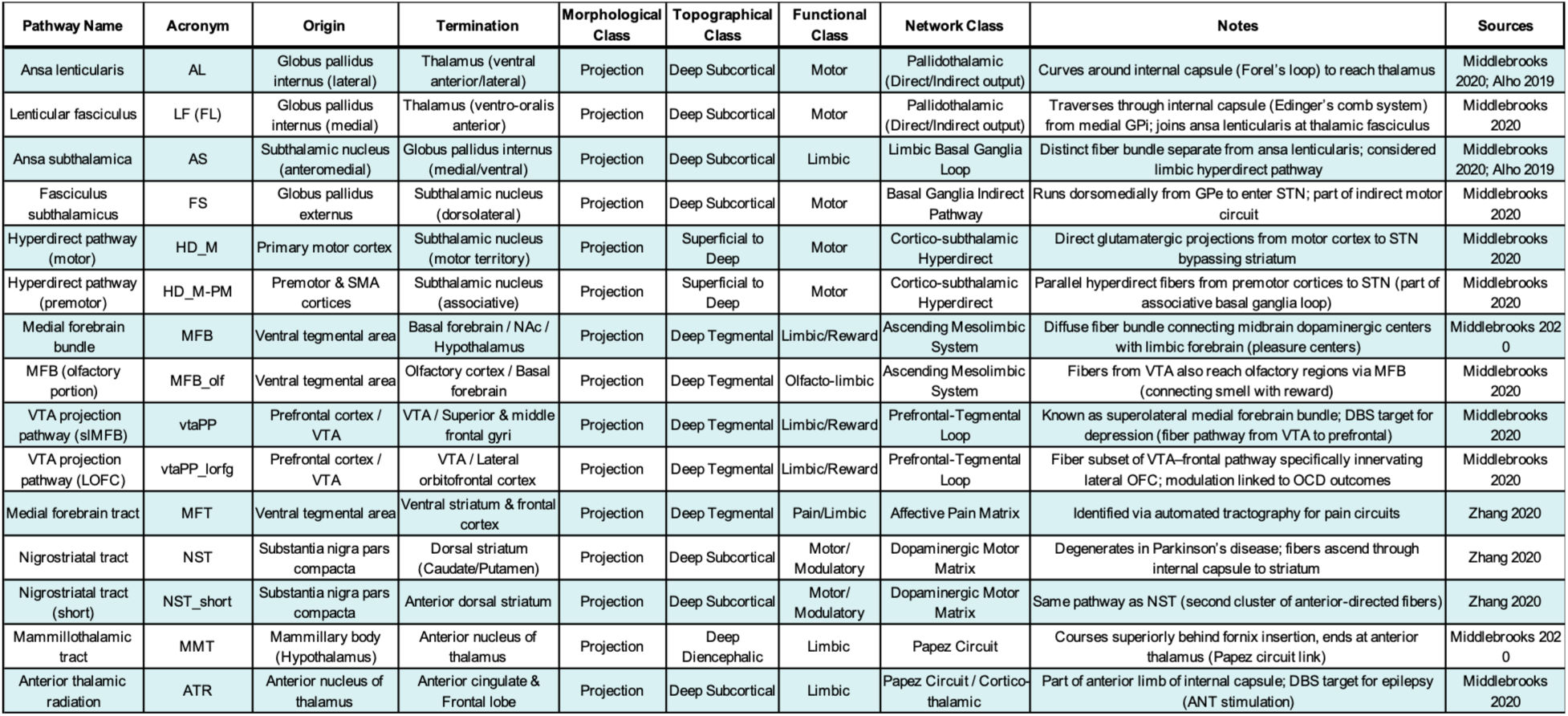
Basal Ganglia, Limbic, and Reward Pathways.

Table 2 contains the cerebellar peduncular systems and their major rubral, thalamic, and cortical continuations. These pathways represent the principal inflow and outflow systems of the cerebellum, including superior, middle, and inferior peduncular components as well as dentatorubrothalamic and related cerebellar output trajectories.

**Table 2.**
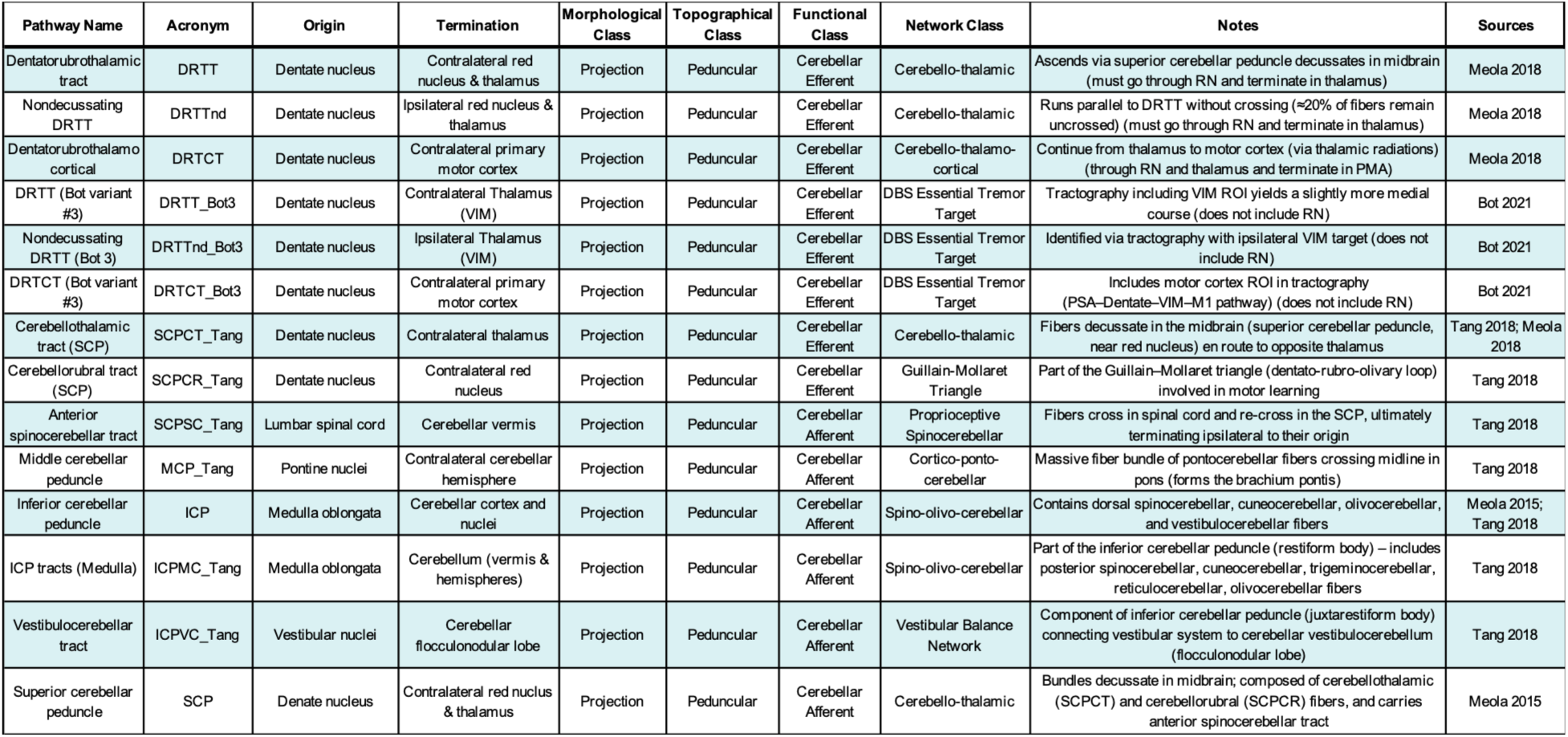
Cerebellar Peduncular Systems.

Table 3 contains the major superficial sensorimotor projection pathways of the ventrolateral brainstem. These include descending motor systems such as the corticospinal and corticopontine pathways, together with ascending sensory relays such as the medial lemniscus, spinothalamic tract, and lateral lemniscus.

**Table 3.**
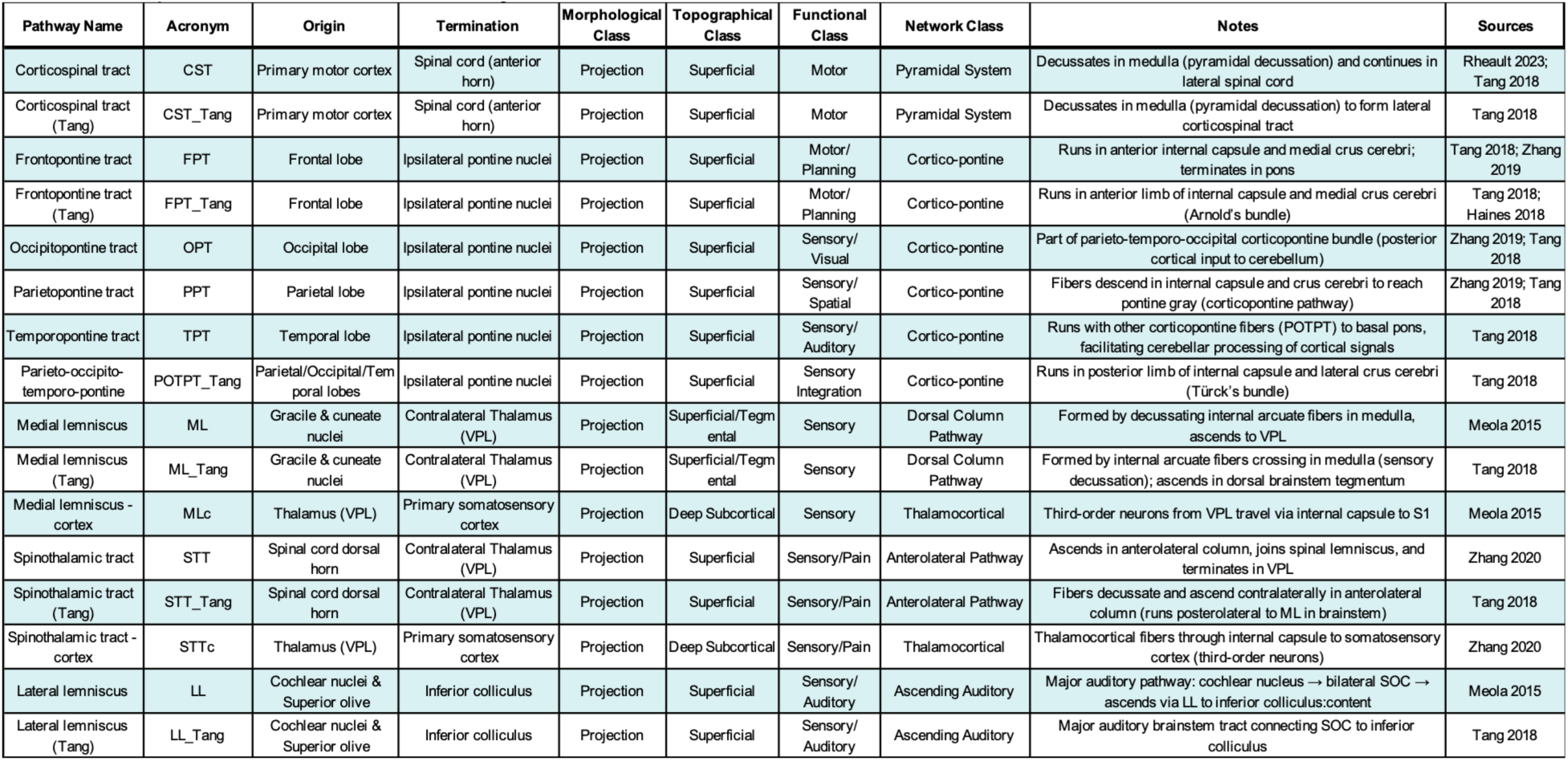
Superficial Sensorimotor Projections.

Table 4 contains the deep intrinsic and homeostatic pathways of the tegmental core. These pathways remain within, or closely tied to, the deep brainstem and include pathways involved in local motor coordination, oculomotor control, autonomic regulation, postural tone, and descending homeostatic modulation.

**Table 4.**
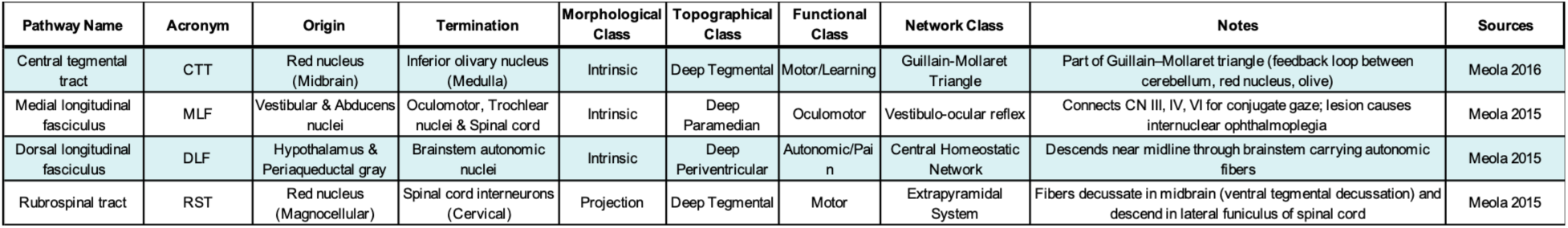
Deep Intrinsic and Homeostatic Pathways.

#### 2.3.3. Notes on terminology, biological interpretation, and variants

Because we are reconstructing a highly diverse set of neuroanatomical structures, it is important to clarify the terminology and biological interpretation of the resulting outputs. We use the following terms:

- Tract: A specific, continuous, and typically monosynaptic axonal projection linking two anatomical regions (e.g., the corticospinal tract).
- Pathway / Route: A broader functional-anatomical trajectory that may involve multiple synaptic relays along its course.
- System / Circuit: A large-scale functional grouping of interrelated tracts and pathways.
- Bundle: The tractography-derived object or physical segmentation mask produced by our automated pipeline to represent a tract or pathway in image space.

Crucially, while many of our segmented bundles represent direct monosynaptic tracts, others are continuous tractographic representations of large-scale, polysynaptic connectivity systems. In the anatomical, tractographic, and neuromodulation literature, these pathways often have overlapping or multiple definitions. When multiple definitions were tractographically reproducible, we did not attempt to endorse a single anatomical interpretation; rather, we retained the definitions used in the source references to maximize the utility of the framework.

This biological distinction is particularly relevant for the dentatorubrothalamic system. Classical definitions of the dentatorubrothalamic tract require passage through the red nucleus, where others isolate a direct dentate-to-thalamic trajectory [24]. We therefore retained both the classical DRTT-style definitions and “Bot-2021” [24] variants (as well as nondecussating routes). Furthermore, we explicitly include the dentatorubrothalamocortical tract (DRTCT), which represents the cerebello-thalamo-cortical continuation. From a neuroanatomical perspective, the DRTCT is inherently polysynaptic. Because it is reconstructed and learned by the network as a single continuous object, it should be interpreted as a standardized tractographic representation of a larger functional-anatomical route, rather than direct evidence of a single monosynaptic axonal pathway.

A similar interpretation applies to the ventral tegmental and forebrain reward pathways. The superolateral medial forebrain bundle, the ventral tegmental area projection pathway, and related orbitofrontal variants are not used consistently in the literature [11, 23, 37], with reconstructions overlapping with or partially reflecting other frontal-subcortical systems. We retained the names and definitions used in the cited references without making claims about preferred nomenclature. The medial forebrain tract was also kept as a distinct pathway because it has been described separately in tractographic studies in work related to pain and limbic circuitry [20, 38, 49].

We also retained the distinction between the ansa lenticularis and ansa subthalamica. Although historically treated as related or partially overlapping pallidofugal systems, more recent anatomical and imaging work supports their separation on the basis of trajectory and connectional anatomy [25].

Some pathways with the same nominal origin and termination separated into more than one reproducible tractography cluster across subjects (e.g., in vtaPP, NST, and MFB). We retained cluster-specific variants when the separation was stable enough to support consistent labeling. These distinctions should be interpreted as tractography-derived subdivisions and not necessarily as definitive anatomical subdivisions established by histology or dissection.

Pathways labeled with the suffix _Tang were derived from the probabilistic brainstem atlas of Tang and colleagues [16]. These bundles generally represent the brainstem portion of larger projection systems and do not necessarily attempt to reconstruct the full cortical continuation of the pathway. Conversely, thalamocortical continuations were explicitly included (e.g., MLc, STTc) when representing the secondary relays of their respective sensory systems.

Ultimately, all outputs produced by this framework are best understood as reproducible, anatomically informed, tractography-derived representations of connectivity systems curated for quantitative analysis and surgical targeting, grounded in their respective operational definitions.

### 2.4. Reference label generation

#### 2.4.1. Reference bundle masks and parcellations

For each retained pathway from each HCP subject, the tractogram was processed into a voxelwise bundle mask and ordered along-tract profile labels using the same bundle-to-label strategy used to create the original BundleParc training targets [30]. A population-averaged centerline was computed for each pathway from streamlines transformed into MNI space using ANTs nonlinear registration. The centerline was mapped into each subject’s space and used to establish consistent streamline orientation (i.e., left-to-right, posterior-to-anterior, inferior-to-superior) and along-tract correspondence. Bundle masks and parcel labels were generated from each subject’s tractography. The pathway was then divided into 50 ordered segments from origin to termination using a hyperplane-based procedure [50]. Briefly, a subset of streamlines was resampled along bundle length, maximally separating hyperplanes between adjacent along-tract positions were estimated with a support vector machine, and voxels within the binary bundle mask were assigned parcel identities by classifier prediction. The resulting parcel maps ranged from 1 to 50 within the bundle and 0 outside the bundle. After label generation in subject space, parcel values were normalized to [0,1], with 0 reserved for background, and smoothed within the bundle mask using Gaussian normalized convolution. This yielded continuous label maps that increased monotonically from pathway origin to termination. Binary reference masks were defined by the nonzero values the corresponding label maps.

#### 2.4.2. Preprocessing into model space

All reference masks and parcellation maps were defined in a 1 mm isotropic resolution space, and cropped to a fixed 176 × 176 × 176 field of view without altering aspect ratio. For each subject, four fODF representations were computed in the same model space to expose the network to variation in angular sampling, response estimation, and tissue modeling [51]. These included multi-shell multi-tissue CSD using the full HCP diffusion acquisition, single-shell CSD using only 12 directions from the b = 1000 shell, single-shell single-tissue CSD using the full b = 1000 shell, and multi-shell multi-tissue CSD using the full b = 1000/2000/3000 shells in the scilpy implementation (with a fixed tensor response function). The resulting fODF volumes and reference labels were stored on the same 1 mm isotropic cropped grid and formed the paired inputs and supervision targets used for network training, validation, and testing.

### 2.5. BundleParc segmentation network

#### 2.5.1. Network Architecture

We adopted the published BundleParc [30] framework and retrained it on the brainstem and subcortical pathway set. Briefly, BundleParc is a 3D encoder-decoder network based on a U-Net-style architecture with skip connections and two-way cross-attention blocks in the decoder. The network takes as input a cropped, 1 mm isotropic, spherical harmonics-encoded fODF volume together with a one-hot prompt specifying the pathway to be predicted. For each requested pathway, the network outputs a binary bundle mask and a continuous along-tract parcellation map. A single shared network was used for all pathways. **Figure 2** summarizes the architecture, and we point the reader to [30] for detailed description of network architecture.

**Figure 2.**
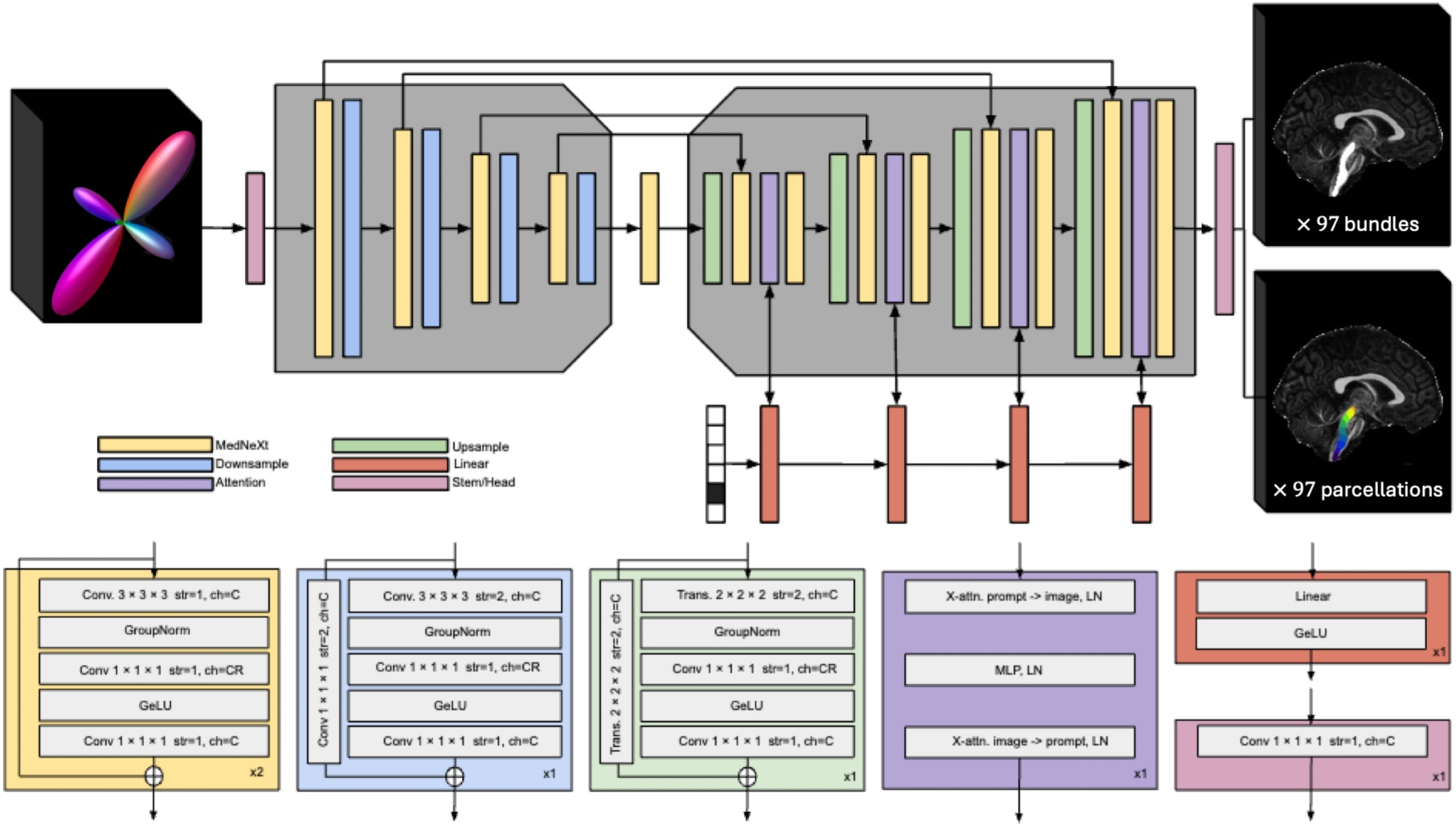
BundleParc segmentation network. The network is based on a U-Net-style encoder-decoder architecture with skip connections and two-way cross-attention blocks in the decoder. Inputs consist of spherical harmonics-encoded fODF volumes together with a one-hot vector indicating the pathway to be predicted. For each requested pathway, the network outputs two volumes, namely a binary bundle mask and a continuous along-tract parcellation map. Conv indicates convolution, str indicates stride, ch indicates number of output channels, Trans indicates transposed convolution, X-attn indicates cross-attention, LN indicates LayerNorm, and MLP indicates multilayer perceptron. The symbol ⊕ denotes addition of parallel outputs, ×2 indicates repetition of the layer, C indicates the number of output channels, and CR indicates the number of output channels multiplied by the expansion ratio. Adapted from the original BundleParc framework.

#### 2.5.2. Training strategy

Subjects were assigned at the subject level to training, validation, and test splits of 840, 105, and 105 subjects, respectively. Because the number of retained examples varied across pathways after quality assurance, split assignment was performed pseudo-randomly to preserve approximately even pathway-specific coverage across the three subsets.

To improve robustness across acquisition quality and reconstruction assumptions, each training batch used one of the four precomputed fODF representations sampled at random for the selected subject. In this way, the network was exposed to the same reference anatomy under different angular sampling schemes, tissue models, and response-function assumptions. This strategy was inspired by TractSeg [51], which sampled three peak representations during training, and extended here to four fODF representations spanning full multi-shell acquisitions, reduced angular sampling, and different reconstruction assumptions.

Following the BundleParc training framework, on-the-fly augmentation included random spatial perturbations and signal perturbations of the input fODFs. These included random translations and rotations, zoom, low-resolution simulation, additive Gaussian noise, and truncation of higher-order spherical harmonic coefficients. Augmentation followed the original BundleParc implementation [30]). Optimization followed the original BundleParc joint loss formulation for simultaneous mask and parcellation prediction. The model was implemented in PyTorch and trained with a batch size of 2 on an NVIDIA RTX 6000 Ada GPU with 48 GB memory. Training proceeded for 50 epochs, with epoch 15 (lowest validation loss) chosen for all subsequent analysis.

### 2.6. Model inference and output generation

For inference on new datasets, we first derived a subject-specific brain mask. We generated a conservative brain, creating the union of a diffusion-derived mask (*dwi2mask* in MRtrix3 [32]) and mask derived from the mean b=0 image (*bet*, FSL [52]). Diffusion volumes were then cropped to the bounding box (with one-voxel padding margin) and resampled to 1mm isotropic resolution. Response function estimation and fODF generation were performed automatically depending on diffusion acquisition. For multi-shell acquisitions, response estimation used multi-shell multi-tissue response and spherical deconvolution (*dwi2response dhollander*; *dwi2fod msmt_csd*). For single-shell acquisitions, response function is estimated using an iterative algorithm for single-fiber selection and estimation[53], with fiber orientations computed using single-shell single-tissue CSD [54]. All inference was performed in this cropped 1 mm isotropic model space.

The trained network was then applied to the subject-specific fODF volume to predict, for each pathway, a binary bundle mask and a continuous along-tract parcellation map. We used N=10 parcels for all analyses as an example of the discretization we would select for a typical along-tract analysis. The number of parcels is user-defined and can be changed at inference without retraining the model. Predicted outputs were written in the common cropped model space and were used for quantitative evaluation. When streamline representations were desired, these outputs were subsequently used for the optional tractography procedure (described in Streamline Reconstruction below).

The end-to-end processing, inference, and tractography is available as a Singularity container [55], source code, and an executable trained model to facilitate evaluation (See *Data & Code Availability*).

### 2.7. Quantitative evaluation

#### 2.7.1. Held-out test set performance

Performance was evaluated on held-out HCP test subjects by comparing model outputs with the tractography-derived reference labels. Binary bundle masks were evaluated with the Sørensen– Dice index, which measures voxelwise overlap between the predicted and reference masks (ranging from 0 to 1, with 1 indicating perfect overlap). Along-bundle parcellations were evaluated with mean absolute error (MAE) between predicted and reference parcel labels, computed only within voxels shared by both masks (indicating, on average, how similar the labels are). Metrics were computed for each subject-pathway pair and summarized across pathways. We also examined associations between pathway size and Dice/MAE across the 97 pathway labels using Spearman correlation.

#### 2.7.2. External generalizability

To assess out-of-distribution generalizability, the trained model was applied to the 18 external datasets described above. For each dataset, 10 subjects were selected at random, and all predicted pathways were reviewed visually as left-right paired parcellations. Although this is a small sample within each individual cohort, it yielded 180 subjects and 17,460 predicted subject-pathway outputs for exhaustive quality assurance [56]. Three team members trained to identify tractography failures (K.G.S., M.E.K., and G.R.) performed visual quality assurance across the full pathway set. Dataset and diagnostic information were not displayed during review. Predicted pathways were assigned pass or fail labels based on anatomical plausibility, including origin, termination, spatial location, trajectory, and completeness of ordered parcellation. Results were summarized as pass rates by pathway and by dataset.

### 2.8. Optional streamline reconstruction from predicted segmentations

The primary output of the framework is a subject-specific bundle segmentation and ordered parcellation in image space. In applications where streamline representations may be desired we make available a tractography procedure constrained by the predicted labels. For each pathway, tractography was run in MRtrix3 from the subject-specific white matter fODF image using *tckgen* with iFOD2 propagation – with seeding restricted to the bundle mask (--seed), segmentation was used as the tracking mask (--mask), and the first and last parcel labels were used as inclusion regions (--include) to enforce passage from pathway origin to termination.

This approach provides an anatomically constrained mechanism for generating streamline representations from the predicted parcellations [46]. We note, however, that this step was intended as a practical downstream option rather than an optimized tractography protocol, as many alternative choices for seeding, propagation, filtering, and endpoint constraints are possible given a bundle parcellation.

## 3. Results

### 3.1. Bundle Segmentation

The trained network generates subject-specific bundle segmentations and ordered along-bundle parcellations across the full pathway set. Figure 3 shows representative predictions from a held-out test subject. Across systems, predicted bundle parcellations follow expected anatomical location and trajectory, and parcel labels progress smoothly from one end of the pathway to the other.

**Figure 3.**
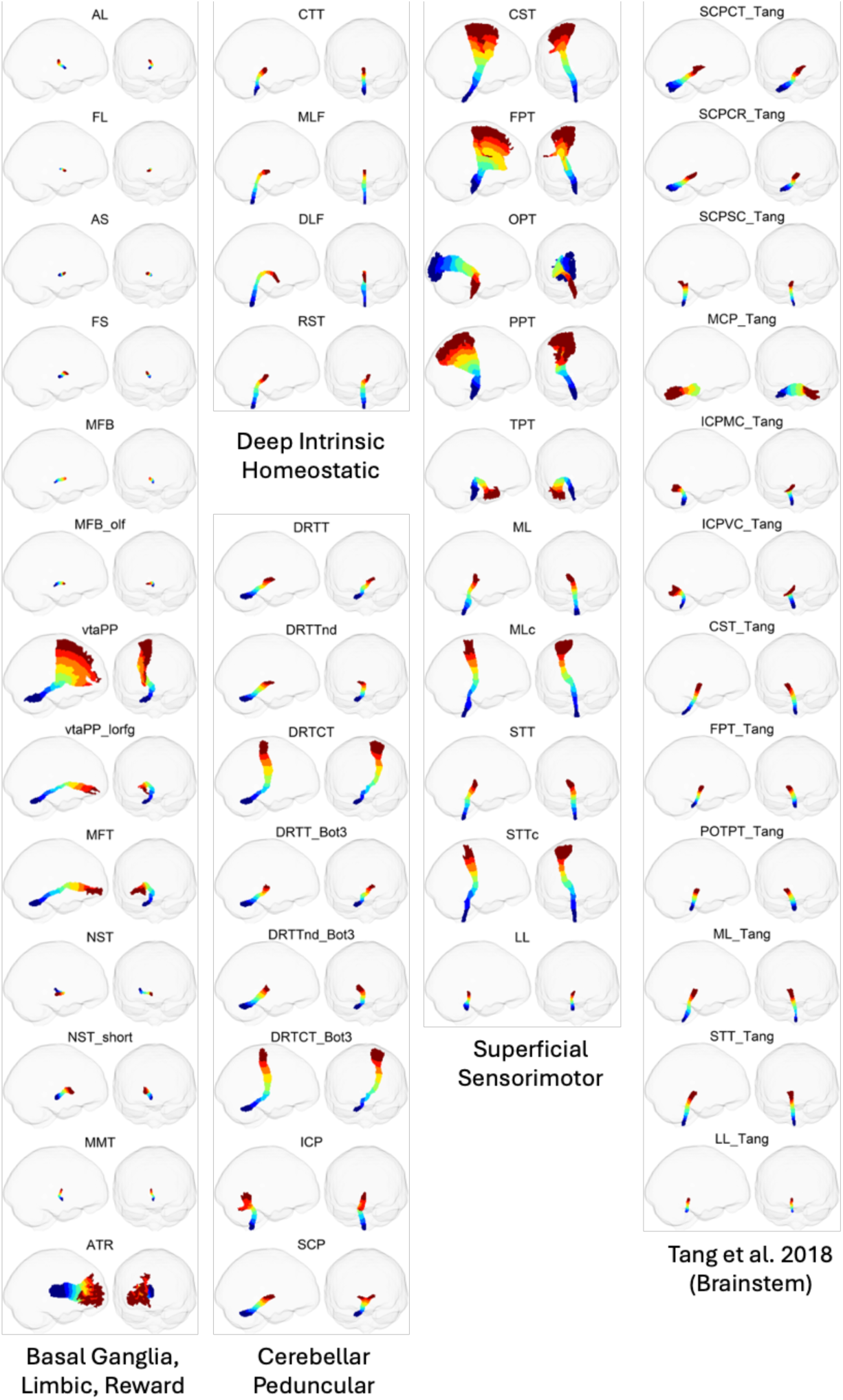
Representative bundle segmentations and parcellations. Predicted pathways are shown as subject-specific parcellations in image space for one randomly selected test subject not used for training or validation. For each pathway, sagittal and coronal views are shown within a glass-brain. Parcel labels are displayed from low to high (values 1-10) as a blue-to-red progression. Pathways are arranged according to the major anatomical groups summarized in Tables 1–4, with Tang-derived brainstem pathways shown in a separate column.

Predicted pathways span a broad range of sizes and extents. Shorter bundles (including the ansa lenticularis, lenticular fasciculus, ansa subthalamica, fasciculus subthalamicus, medial forebrain bundle, and mammillothalamic tract) are segmented as compact structures within the deep subcortical and brainstem regions. Longer bundles (including corticospinal, corticopontine, ventral tegmental, and thalamocortical pathways) extend over larger anatomical territories while maintaining ordered parcellation from origin to termination. Within the brainstem, several pathways displayed similar overall trajectories but remained spatially distinct by their anterior-posterior or medial-lateral offsets, as seen in the deep intrinsic tracts, major ascending sensorimotor systems, and Tang et al. pathways.

A useful feature of the predictions is that ordered parcellation remained anatomically coherent across both short and long pathways, always from left-to-right, posterior-to-anterior, or inferior-to-superior. For shorter trajectories, the smaller spatial extent reduces separation between parcels, and simple binary segmentation may be preferred for quantitative analysis.

### 3.2. Held-out test performance

On the held-out HCP test set, binary segmentation performance is moderate to high across most pathways (**Figure 4**). Of the 97 pathways, 89 have an average Dice coefficient of 0.6 or greater. The lowest average Dice values occur in MFB, LL_Tang, AL, vtaPP_lorfg, FL, MMT, FS, and MFB_olf, reflecting a combination of short pathway length, compact anatomy, and complex local geometry. Overall, these Dice values are similar to those reported previously for automated segmentation of larger cerebral white matter pathways [30, 51].

**Figure 4.**
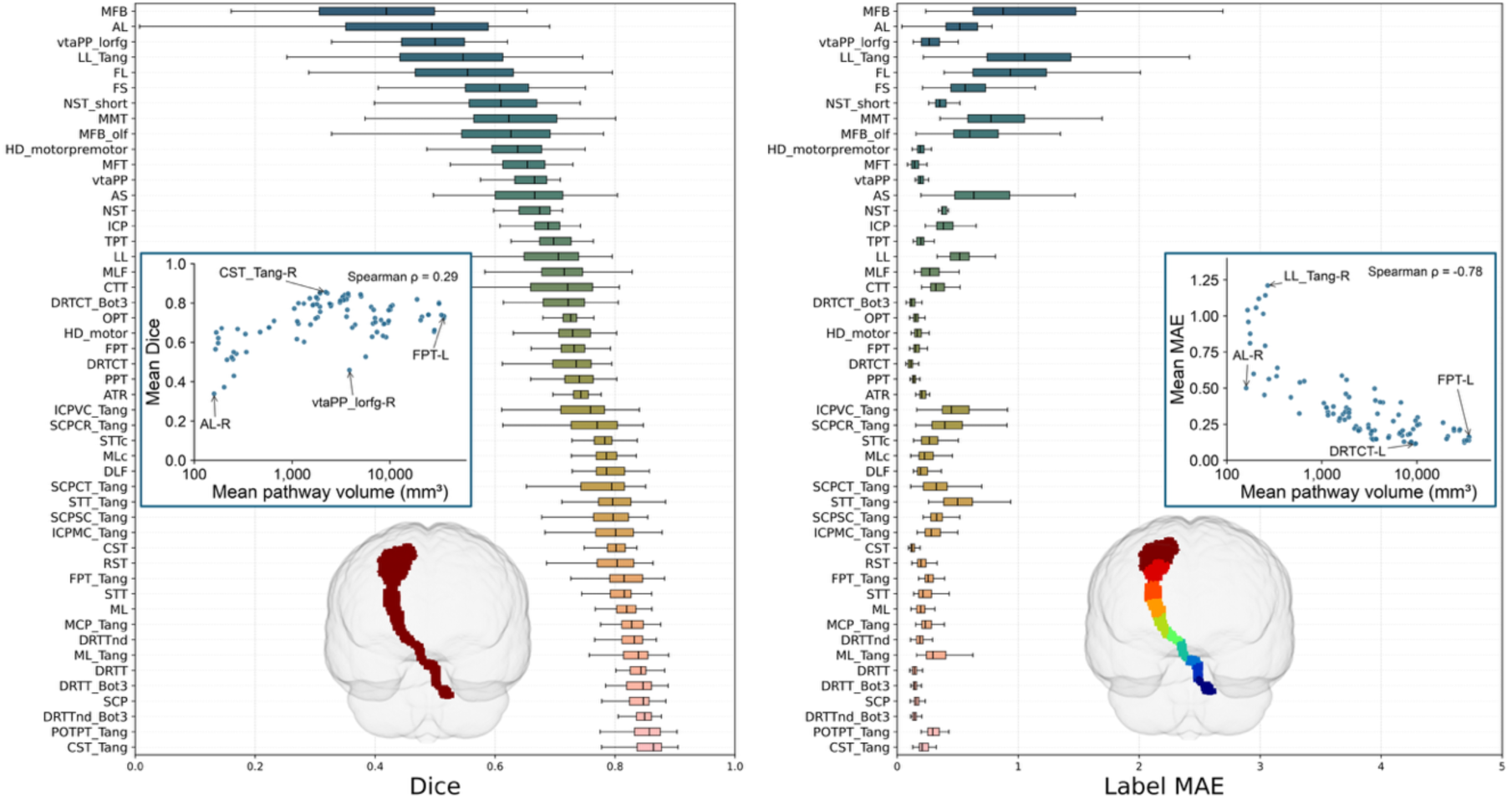
Quantitative performance on the held-out HCP test set. Binary segmentation and parcellation accuracy are summarized across the 105 held-out test subjects. Left, pathways are ranked by average Dice coefficient for binary bundle segmentation. Right, parcellation accuracy is shown as the mean absolute error between predicted and reference parcel labels. Boxplots summarize pathway-wise performance across subjects. Insets show mean Dice/MAE plotted against mean pathway volume. The glass-brain renderings illustrate the two exemplar output representations used for evaluation, binary segmentation for Dice analysis and full ordered parcellation for label error analysis.

Parcellation accuracy is similarly stable across the held-out cohort. Mean absolute error remains low across pathways, with predicted labels on average less than one parcel absolute error from the reference label for most pathways. Pathways with lower binary Dice generally also show larger parcellation error. Larger reference bundles tended to have higher Dice (Spearman ρ = 0.29) and lower parcellation MAE (ρ = −0.78; **Figure 4, insets**). Together, these results indicate that the model preserves both binary pathway extent and ordered parcel structure on unseen test data.

### 3.3. Generalizability across external datasets

We apply the trained model to 18 external datasets spanning multiple sites, age ranges, diagnostic groups, and diffusion MRI acquisition protocols. **Figure 5** shows representative pathway predictions for each dataset, together with dataset name, age range, cohort, and diffusion acquisition characteristics. Across cohorts, data range from infancy to late aging (0-100 years old), include cognitively normal and clinical populations, and vary substantially in spatial resolution, field of view, b value, number of diffusion directions, and single-shell versus multi-shell design.

**Figure 5.**
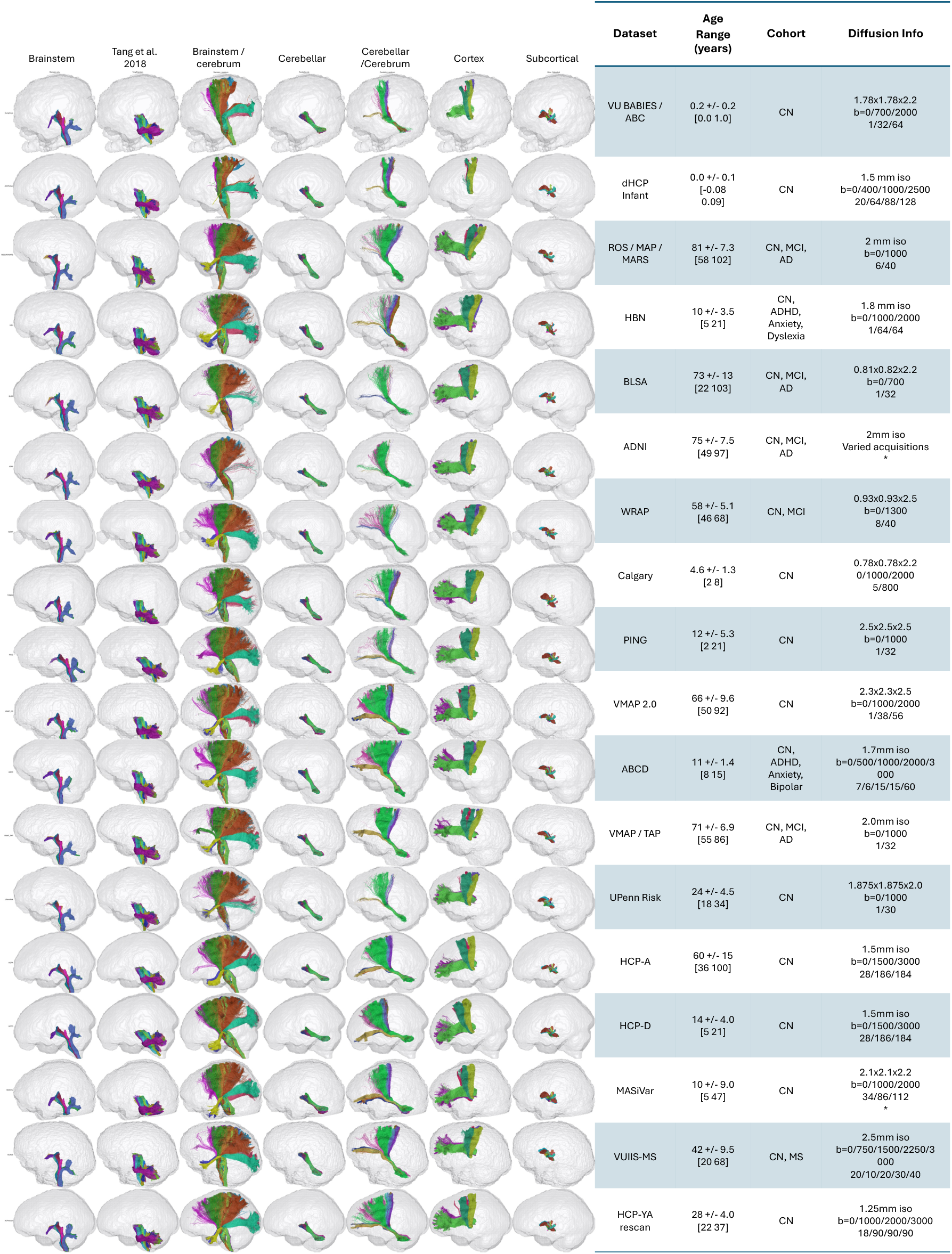
Generalizability across external datasets. Representative bundle segmentations and parcellations are shown for one subject from each of 18 external datasets. For display, pathways are grouped by gross location and termination pattern and visualized as parcellations within a glass-brain. Each panel includes the dataset name, age range, cohort, and diffusion MRI acquisition characteristics, including spatial resolution, b values, and number of diffusion directions. Within each dataset, the displayed subject was randomly selected from those that passed visual quality assurance. Cohort abbreviations: CN for cognitively normal, MCI for mild cognitive impairment, AD for Alzheimer disease, ADHD for attention-deficit/hyperactivity disorder, and MS for multiple sclerosis. Asterisk (*) indicates varied acquisition parameters.

Predictions remain anatomically plausible across most pediatric, young adult, and aging datasets despite dataset heterogeneity. **Figure 6** summarizes pathway success rates by dataset and by bundle. The most consistent failures occurred in the two infant datasets, where predictions were poor across much of the pathway set. Outside of infancy, most datasets show high success rates, with several cohorts approaching complete recovery of most bundles. Others, including ROSMAPMARS, HBN, and BLSA, typically fall in the 80–90% success range. Across pathways, vtaPP_lorfg and MMT fail most consistently, and ICP also shows reduced success relative to most other bundles.

**Figure 6.**
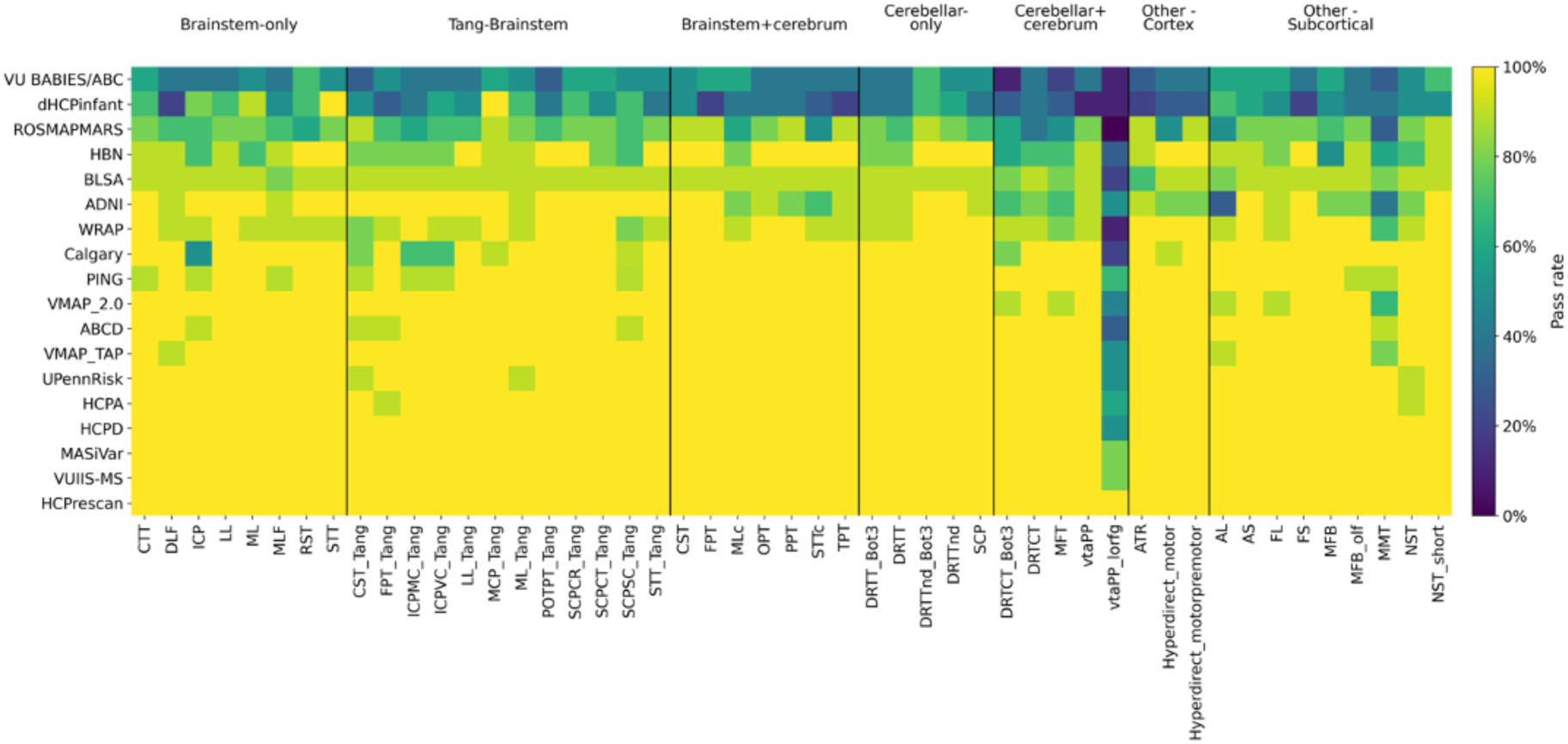
Quality assurance across external datasets and pathways. Success rates are summarized by dataset (rows) and pathways (columns) as the proportion of predicted bundles judged anatomically plausible on visual quality assurance. Quality assurance was performed on a pass or fail basis for every pathway in every subject [56], considering pathway origin, termination, spatial location, trajectory, and completeness of ordered parcellation.

Failures are not random and generally reflect identifiable properties of the input data. The most common failure mode is incomplete parcellation, in which one or both terminal parcel labels are not fully recovered. In some subjects this is associated with a limited field of view, particularly when substantial portions of the inferior brainstem or cerebellum are not within the acquired field of view. This is most evident in datasets such as ROSMAPMARS and HBN, where incomplete coverage can affect both brainstem and non-brainstem pathways. Other failures are associated with severe motion artifact or markedly oblique acquisition geometry (whereas training data is largely AC-PC oriented, with train-time augmentations). Of note, one BLSA subject fails across nearly the full pathway set because of pronounced volume misalignment (we elected not to exclude any randomly selected subjects from analysis).

### 3.4. Tractography across representative systems

To illustrate the range of pathway systems captured by the framework, we show the automated tractography representations from six representative pathway groupings (**Figure 7**; single-shell VMAP/TAP examples in **Supplementary Figure 1**). These include the Tang brainstem atlas set, the basal ganglia and hyperdirect pathways, the dentatorubrothalamic neuromodulation set, the ventral tegmental reward network, the affective and somatosensory pain-related pathways, and the deep intrinsic brainstem tracts. These examples highlight the ability to resolve unique pathways, and also the short, compact, and sometimes highly overlapping pathways within the subcortex and brainstem.

**Figure 7.**
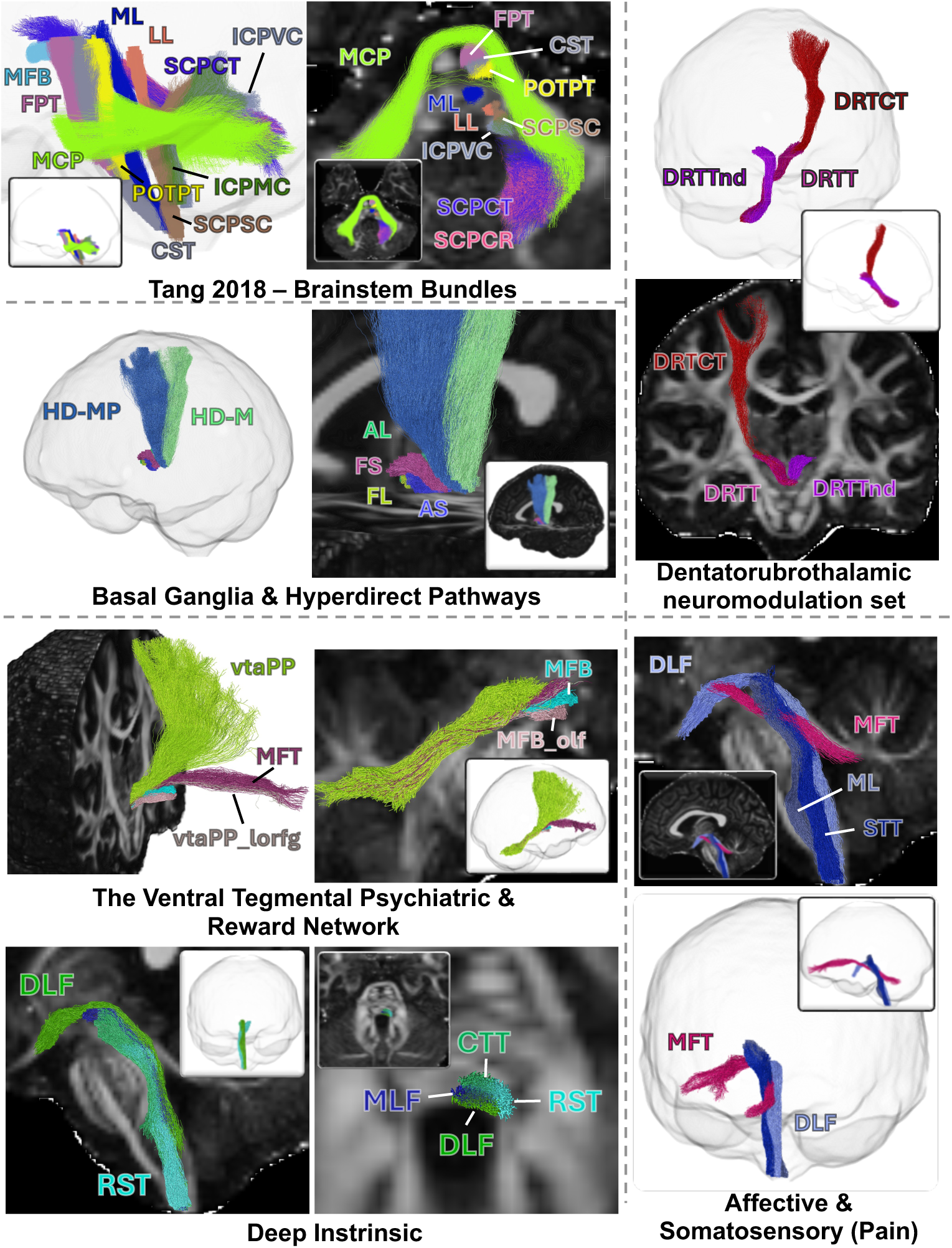
Representative streamline reconstructions across six pathway systems. Streamline representations generated from predicted segmentations are shown for six pathway groupings (in a held-out HCP subject) selected to illustrate the range of anatomy captured by the framework. Distinct colors indicate different pathways within each grouping. Panels include Tang-derived brainstem pathways, basal ganglia and hyperdirect pathways, dentatorubrothalamic neuromodulation pathways, ventral tegmental reward pathways, affective and somatosensory pathways, and deep intrinsic brainstem tracts. Views include whole-brain renderings, sectional views, and zoomed insets to highlight local organization and overlap. Supplementary Figure 1 shows corresponding examples from a single-shell VMAP/TAP acquisition.

Within the “Tang” [16] brainstem pathways, the major ventral projection systems (frontopontine, corticospinal, and parieto-occipito-temporo-pontine pathways) remain clearly separated, traveling in parallel through the ventral brainstem while maintaining distinct topological positions. A similar organization is visible among the major ascending and peduncular pathways, including the medial lemniscus, lateral lemniscus, spinocerebellar, vestibulocerebellar, and superior cerebellar peduncular components, where neighboring bundles follow closely related trajectories but remain individually identifiable.

The dentatorubrothalamic group highlights the structural differences between decussating and nondecussating cerebellothalamic pathways. The classical DRTT and DRTTnd separate clearly at the level of the superior cerebellar peduncle and midbrain, while DRTCT extends further rostrally as a cerebello-thalamo-cortical continuation. Their trajectories show the ability of the algorithm to resolve the complex crossing geometry of the mesencephalic region.

The basal ganglia and hyperdirect pathways show the compact geometry of the subthalamic region. The ansa lenticularis, lenticular fasciculus, fasciculus subthalamicus, and ansa subthalamica occupy closely neighboring trajectories with substantial local overlap, while the motor and motor-premotor hyperdirect pathways diverge toward different cortical territories. These streamlines emphasize the dense arrangement of overlapping tracts that are primary targets for Parkinson’s disease treatments.

The ventral tegmental grouping captures a predominantly anterior-posterior system linking the midbrain with ventral forebrain and frontal regions. One can appreciate the slight difference in terminations in the two MFB bundles (MFB, MFB_olf), and strong similarities between MFT and vtaPP_lorfg (which have been similarly described in the literature).

The affective and somatosensory grouping highlights the spatial relationships between ascending sensory and pain-related pathways. The medial lemniscus and spinothalamic tract maintain distinct but neighboring courses through the brainstem, while the dorsal longitudinal fasciculus occupies a more dorsal and midline position and the medial forebrain tract traverses the same region along a different axis. The resulting streamlines show both the anterior-posterior and medial-lateral organization of these pathways as they pass through the compact tegmental core.

The deep intrinsic set demonstrates that even the smallest tegmental pathways can be reconstructed as distinct streamline bundles. The central tegmental tract is positioned relatively anteriorly, the medial longitudinal fasciculus remains immediately adjacent to the midline, the rubrospinal tract follows a more lateral course, and the dorsal longitudinal fasciculus occupies a more dorsal position. Although these pathways lie in close proximity and overlap in projection, their relative positions are preserved across the length of the brainstem.

## 4. Discussion

We present an automated approach for segmentation and along-tract parcellation of 97 subcortical and brainstem white matter pathways. Historically, the dense and complex architecture of the deep tegmental core, basal ganglia, and cerebellar circuits has rendered these regions difficult to study with conventional in vivo diffusion MRI, leaving them underrepresented in large-scale connectomics and brain mapping analyses. By combining curated anatomical pathway definitions, HCP diffusion MRI data, and a BundleParc-based segmentation model, our method enables automated extraction of subject-specific bundle masks and ordered parcellations across a broad set of white matter systems. This enables the segmentation and study of these pathways across datasets that vary in age, disease context, acquisition protocol, and image quality, including clinically feasible diffusion MRI acquisitions.

### 4.1. Extending automated tract segmentation to deep brainstem and subcortical pathways

An important motivator of the present study is recent ultra-high-resolution ex vivo imaging by Maffei and colleagues [14] demonstrating that the intricate subcortical connectome can be mapped in fine anatomical detail when image resolution and signal-to-noise ratio are sufficiently high. Their work provided a proof of concept that these networks are tractographically recoverable, especially when well-defined anatomical rules (i.e., where pathways start, stop, and do not go) are used to inform the tractography process [46]. Our approach follows the same general principle, we use curated pathway definitions, with the goal to train a model that can recover these systems in new subjects and in conventional diffusion MRI datasets.

The closest automated precedent is the BrainStem Bundle Tool developed by Olchanyi et al. [26], which used ex vivo diffusion MRI, histology-validated probabilistic fiber maps, and a convolutional neural network to segment eight bilateral rostral brainstem white matter bundles. Their results showed that automated segmentation of these small bundles is feasible across in vivo and ex vivo datasets, and that tract-resolved brainstem measures can reveal disease-related alterations in neurodegenerative diseases and injury. Our work extends this direction to a broader set of brainstem and subcortical pathways and adds along-tract parcellations for downstream quantitative analysis.

From a practical standpoint, the primary contribution of this framework lies not in the underlying deep learning architecture, but in the extensive anatomical curation required to train it. The vast majority of our effort was dedicated to generating reliable reference streamlines for all pathways. This process required iteratively refining tractography protocols, defining start and end regions, refining inclusion and exclusion rules, iterating over tractography failures, and manually reviewing the resulting bundles across subjects. The final product (i.e., the trained model) is valuable because it can execute these highly specialized segmentations on new, unseen datasets automatically, and in a few minutes.

This work also builds on the broader movement toward direct tract segmentation. TractSeg [51] (as well as other approaches, including Tracula [5]) demonstrated that white matter bundles can be segmented directly from fiber orientation information, without requiring tractography (i.e., propogating streamlines), atlas registration, or cortical parcellation during inference. Recently, BundleParc extended this concept to ordered bundle parcellation, producing tractometry-ready label maps directly from fiber orientation distribution volumes and avoiding several sources of variability introduced by streamline reconstruction and centerline-based parcellation. These features are particularly well suited to deep brainstem and subcortical anatomy, where small changes in tractography parameters, endpoint rules, or registration quality can have large effects on the apparent location and extent of a pathway.

### 4.2. Generalizability and segmentation robustness

To be scientifically and (potentially) clinically useful, an automated segmentation tool must perform reliably on data outside its training distribution. When applied to external datasets spanning pediatric, aging, and neurodegenerative cohorts, our framework successfully recovered these subcortical pathways despite substantial variations in spatial resolution, angular sampling, and disease state.

However, we observed specific failure modes that users must account for in their experimental design. Frequent failures in the evaluated infant datasets indicate that the current model is not immediately suitable for those data without further adaptation and validation. Most failures involved incomplete label sets, with missing cortical extensions or brainstem segments that varied across subjects, with no clear pattern suggesting consistent regional locations or particular pathway groups. Failure is likely because unmyelinated tissue properties and rapidly developing anatomy diverge too far from the adult training distribution. While these findings do not establish a precise age threshold for reliable performance, we recommend critical assessment of results if used on infant cohorts.

Outside of infancy, failures can occasionally appear random (potentially driven by abnormal local anatomy) but typically stem from identifiable imaging constraints. For example, when the inferior brainstem or cerebellum is truncated by a restricted field of view, or when highly oblique acquisition geometries disrupt the expected spatial priors, the model may generate missing or incomplete parcellations. Incomplete parcellations due to truncated brainstem field of view often still correctly identify the available anatomy, starting (inferiorly) at a mid-range parcellation value (i.e. >1). We also noted that certain complex bundles, such as the vtaPP_lorfg navigating a dense bottleneck above the brainstem, exhibit higher failure rates, and that unexpected left-right asymmetries can occasionally emerge in subcortical systems. As a best practice, we highly recommend the visual inspection of every pathway before conducting downstream analyses. To facilitate this, our released container includes an automated quality assurance module that generates multi-view visual reports for every predicted bundle, allowing users inspect anatomical plausibility.

Finally, we note that for several short bundles, along-tract analysis (e.g., profiling microstructural metrics across ordered segments) may not be anatomically meaningful. Specifically, pathways such as the ansa lenticularis (AL), ansa subthalamica (AS), lenticular fasciculus (FL), fasciculus subthalamicus (FS), medial forebrain bundle (MFB), and nigrostriatal tract (NST) are often shorter than the spatial resolution required to meaningfully separate their intended parcels.

Indeed, many of these structures are better interpreted as compact systems of connections or tractographic representations of functional-anatomical routes, rather than traditional long-range projection tracts with a stable principal axis. For these specific targets, we advocate using the binary segmentations for spatial visualization (e.g., in relation to deep brain stimulation targets) or extracting whole-bundle summary statistics, rather than attempting ordered tractometry. Yet, even when individual parcels are too small for meaningful along-tract analysis, the ordered labels can still define seed and termination regions for automated tractography. The ability to reconstruct these small, anatomically complex pathways automatically is an important practical benefit of the parcellation.

### 4.3. Methodological Considerations

Our primary goal was to generate pathway segmentations and ordered parcellations, rather than streamlines. In many connectomic workflows, streamline representations are ultimately reduced to either binary masks or gridded along-tract coordinates to extract quantitative microstructural metrics. Following the rationale of the BundleParc framework, predicting these end-stage representations directly from fiber orientation distributions circumvents the computational burden and potential errors associated with whole-brain streamline generation and subsequent virtual dissection. Streamlines nevertheless remain useful. They provide an intuitive representation for visualization, surgical context, and comparison with the tractography literature. They may also provide a more spatially specific representation of a pathway than a voxel-wise parcellation alone, especially when the goal is to inspect a trajectory or its relationship to nearby nuclei, electrodes, lesions, or stimulation fields. To accommodate these use cases, we chose to also make available an optional post-inference tractography module. By confining the generated streamlines entirely within the boundaries of the predicted segmentation masks, connecting origin to termination, we ensure the trajectories adhere to the anatomical priors established during the model training phase.

Several methodological choices could be revisited in future work. We selected a specific voxel size, input representation, image field of view, and network configuration, but other choices are possible, including FOD coefficients, FOD peaks, raw diffusion-derived features, different cropped image sizes, and alternative segmentation architectures. A TractSeg-like strategy could also be used to explicitly predict tract orientation maps for bundle-specific tractography, although this was not required for the main goal of segmentation and parcellation. More broadly, automated tract segmentation includes a growing range of atlas-based, clustering-based, tractography-based, hybrid, and deep learning methods. To support future comparisons and alternative training strategies, we make the curated streamlines and derived labels available with the released code and data resources.

### 4.4. Limitations and Future Directions

There are several limitations to this study. First, diffusion MRI (and any medical imaging) will always be limited in the ability to resolve fine structures due to partial volume effects and limited spatial resolution. Second, the reference labels are tractography-derived, not histological ground truth, and are a direct reflection of rules, atlases, and definitions used to create them. Third, although the current pathway set is broad, it is not complete. The framework can be expanded as new anatomical resources become available. An excellent example is a recent ultra-high-resolution subcortical atlas from Friedrich and colleagues [28] which includes over eighty highly specific white matter tracts from ex vivo imaging and histology. Transfer learning could be used from this detailed N=1 sample to new in vivo datasets, or the rules, definitions, and pathways in the atlas could be used to generate reference labels on HCP-like datasets. As noted by Friedrich and team [28], “the Federative Community on Anatomical Terminology lists 455 structures in the subcortex” – motivating the ability to automatically segment and study much more than the currently provided 97 pathways. Fourth, input representations and augmentation choices were not exhaustively evaluated - fODF representations covered a broad range of settings, but not all acquisition and reconstruction combinations; noise was not scaled by SH order; representation and augmentation strategies (including SH rotations implemented with Wigner D matrices) could be evaluated in future work. Finally, future work should also evaluate the framework in specific biological and clinical applications, including development, aging, Parkinson disease and other neuromodulation targets, Alzheimer disease with early brainstem involvement such as locus coeruleus pathology, multiple sclerosis with brainstem or spinal cord lesions, traumatic brain injury, pain, arousal, and autonomic dysfunction.

## 5. Code & Data Availability

The end-to-end preprocessing, inference, quality assurance, and optional tractography workflow is released as a Singularity container available to be downloaded through Zenodo (https://doi.org/10.5281/zenodo.19343084). We also release a young-adult population-averaged atlas and the curated subject-specific reference streamlines and derived labels used for training. The trained model weights are also packaged and available in the container to facilitate independent evaluation and reuse. The codebase used to train the model is available at https://github.com/MASILab/brainstemseg_singularity to facilitate model-finetuning if necessary.

## 6. Appendix A. Pathway names and abbreviations

The 49 pathway names listed below comprise 48 bilateral pairs and one unpaired label for the middle cerebellar peduncle (MCP_Tang), yielding 97 pathway labels.

Ansa lenticularis (AL); Ansa subthalamica (AS); Anterior thalamic radiation (ATR); Corticospinal tract (CST); Central tegmental tract (CTT); Dorsal longitudinal fasciculus (DLF); Decussating dentatorubrothalamic tract (DRTT); Nondecussating dentatorubrothalamic tract (DRTTnd); Dentatorubrothalamocortical tract (DRTCT); Dentatorubrothalamic tract, Bot variant 3 (DRTT_Bot3); Nondecussating dentatorubrothalamic tract, Bot variant 3 (DRTTnd_Bot3); Dentatorubrothalamocortical tract, Bot variant 3 (DRTCT_Bot3); Lenticular fasciculus (FL); Frontopontine tract (FPT); Fasciculus subthalamicus (FS); Hyperdirect pathway, motor (Hyperdirect_motor); Hyperdirect pathway, motor and premotor (Hyperdirect_motorpremotor); Inferior cerebellar peduncle (ICP); Lateral lemniscus (LL); Medial forebrain bundle (MFB); Medial forebrain bundle, olfactory portion (MFB_olf); Medial forebrain tract (MFT); Medial lemniscus (ML); Medial lemniscus to cortex (MLc); Medial longitudinal fasciculus (MLF); Mammillothalamic tract (MMT); Nigrostriatal tract (NST); Short nigrostriatal tract (NST_short); Occipitopontine tract (OPT); Parietopontine tract (PPT); Rubrospinal tract (RST); Superior cerebellar peduncle (SCP); Spinothalamic tract (STT); Spinothalamic tract to cortex (STTc); Temporopontine tract (TPT); VTA projection pathway, superolateral medial forebrain bundle (vtaPP); VTA projection pathway to lateral orbitofrontal gyrus (vtaPP_lorfg);

Corticospinal tract, Tang atlas definition (CST_Tang); Frontopontine tract, Tang atlas definition (FPT_Tang); Inferior cerebellar peduncle tract from medulla oblongata to cerebellum, Tang atlas definition (ICPMC_Tang); Vestibulocerebellar tract of the inferior cerebellar peduncle, Tang atlas definition (ICPVC_Tang); Lateral lemniscus, Tang atlas definition (LL_Tang); Middle cerebellar peduncle, Tang atlas definition (MCP_Tang); Medial lemniscus, Tang atlas definition (ML_Tang); Parieto-occipito-temporo-pontine tract, Tang atlas definition (POTPT_Tang); Cerebellorubral tract of the superior cerebellar peduncle, Tang atlas definition (SCPCR_Tang); Cerebellothalamic tract of the superior cerebellar peduncle, Tang atlas definition (SCPCT_Tang); Anterior spinocerebellar tract of the superior cerebellar peduncle, Tang atlas definition (SCPSC_Tang); Spinothalamic tract, Tang atlas definition (STT_Tang).

## 7. Appendix B. External Datasets

Generalizability was evaluated in 10 randomly selected subjects from each of the following 18 datasets, for a total of 180 subjects. Dataset names follow Figure 5.

| Dataset | Cohort description |
| --- | --- |
| VU<br>BABIES/ABC | Vanderbilt infant cohorts used to assess early postnatal anatomy. |
| dHCP Infant | Neonatal cohort from the Developing Human Connectome Project [57]. |

| **Dataset** | **Cohort description** |
| --- | --- |
| ROS/MAP/MARS | Religious Orders Study, Memory and Aging Project, and Minority Aging Research Study; longitudinal cohorts of aging and cognitive decline [58]. |
| HBN | Healthy Brain Network; children and adolescents with a range of psychiatric and learning profiles [59]. |
| BLSA | Baltimore Longitudinal Study of Aging; a longitudinal adult aging cohort [60]. |
| ADNI | Alzheimer’s Disease Neuroimaging Initiative; participants spanning normal cognition, mild cognitive impairment, and Alzheimer disease [61]. |
| WRAP | Wisconsin Registry for Alzheimer’s Prevention; an adult cohort studying aging and Alzheimer disease risk [62]. |
| Calgary | Calgary Preschool MRI Dataset; a longitudinal study of early childhood brain development [63]. |
| PING | Pediatric Imaging, Neurocognition, and Genetics; a pediatric and adolescent cohort [64]. |
| VMAP 2.0 | Vanderbilt Memory and Aging Project (different acquisition from VMAP/TAP below) [65]. |
| ABCD | Adolescent Brain Cognitive Development Study; a longitudinal pediatric and adolescent cohort [66]. |
| VMAP/TAP | Vanderbilt Memory and Aging Project and Tennessee Alzheimer’s Project samples [65] (clinicaltrials.gov ID: NCT05372172). |
| UPennRisk | University of Pennsylvania cognitive-training and decision-making study in young adults [67]. |
| HCP-A | Human Connectome Project Aging; an adult lifespan cohort [68]. |
| HCP-D | Human Connectome Project Development; a childhood and adolescent cohort [68]. |
| MASiVar | Multisite, multiscanner diffusion MRI dataset including children and adults [69]. |
| VUIIS-MS | Vanderbilt University Institute of Imaging Science cohort including multiple sclerosis and control participants [70]. |
| HCP-YA rescan | Repeat acquisitions from the Human Connectome Project Young Adult cohort [29]. |

## 8. Appendix C. Samples size of reference pathways for training, testing, validation

Counts indicate the number of QA-passed HCP reference labels available for each pathway. The inventory included 1050 HCP subjects with fODF data and 97 pathway labels.

| Pathway | Left | Right | Midline |
| --- | --- | --- | --- |
| AL | 150 | 148 |  |
| AS | 1024 | 1022 |  |
| ATR | 767 | 766 |  |
| CST | 941 | 938 |  |
| CTT | 1023 | 1023 |  |
| DLF | 1027 | 1027 |  |
| DRTT | 981 | 985 |  |
| DRTTnd | 1018 | 1017 |  |
| DRTCT | 1012 | 1012 |  |
| DRTT_Bot3 | 822 | 827 |  |
| DRTTnd_Bot3 | 1008 | 1000 |  |
| DRTCT_Bot3 | 1006 | 995 |  |
| FL | 909 | 908 |  |
| FPT | 1026 | 1023 |  |
| FS | 1022 | 1021 |  |
| Hyperdirect_motor | 1022 | 1022 |  |
| Hyperdirect_motorpremotor | 1020 | 1020 |  |
| ICP | 1026 | 1026 |  |
| LL | 602 | 602 |  |
| MFB | 533 | 533 |  |
| MFB_olf | 1011 | 1010 |  |
| MFT | 911 | 910 |  |
| ML | 1019 | 1017 |  |
| MLc | 1007 | 999 |  |
| MLF | 987 | 987 |  |
| MMT | 1020 | 1020 |  |
| NST | 98 | 98 |  |
| NST_short | 308 | 308 |  |
| OPT | 980 | 978 |  |
| PPT | 1025 | 1025 |  |
| RST | 939 | 939 |  |
| SCP | 992 | 997 |  |
| STT | 1025 | 1025 |  |
| STTc | 1026 | 1023 |  |
| TPT | 693 | 695 |  |
| vtaPP | 260 | 262 |  |
| vtaPP_lorfg | 569 | 571 |  |
| CST_Tang | 976 | 968 |  |
| FPT_Tang | 1000 | 1000 |  |
| ICPMC_Tang | 1012 | 1012 |  |
| ICPVC_Tang | 1008 | 1008 |  |
| LL_Tang | 924 | 924 |  |
| MCP_Tang |  |  | 1032 |
| ML_Tang | 1009 | 1009 |  |
| POTPT_Tang | 1047 | 1047 |  |
| SCPCR_Tang | 1034 | 1034 |  |
| SCPCT_Tang | 1019 | 1019 |  |
| SCPSC_Tang | 949 | 949 |  |
| STT_Tang | 1029 | 1029 |  |

## 9. Supplementary

**Supplementary Figure 1.**
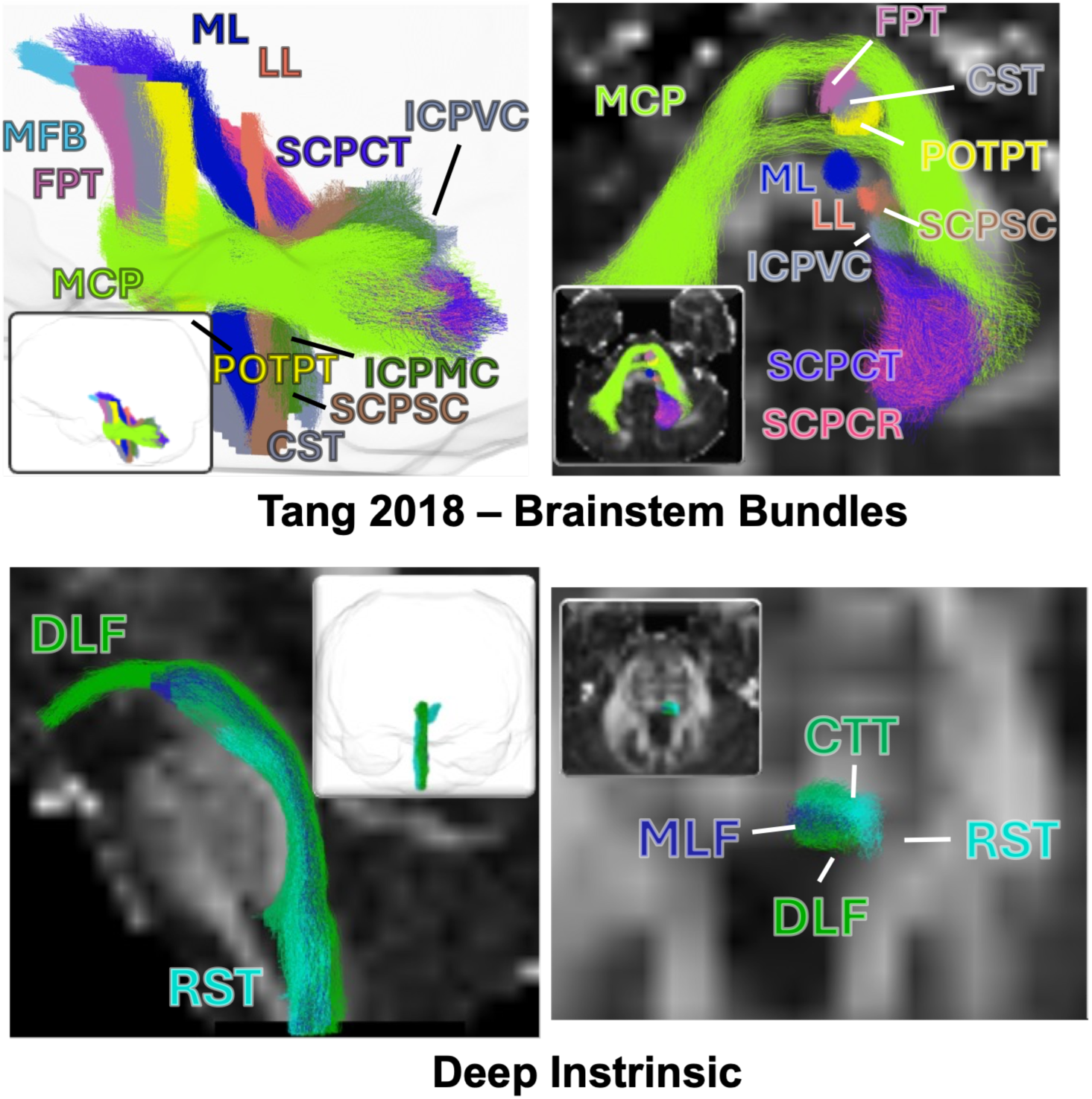
Representative brainstem pathways from a single-shell VMAP/TAP acquisition. Streamline reconstructions generated from predicted segmentations are shown for one VMAP/TAP subject acquired at 2 mm isotropic resolution, with 32 diffusion directions at b = 1000 s/mm². Panels show selected brainstem pathways (top) and the closely packed CTT, RST, MLF, and DLF pathways (bottom), using colors corresponding to Figure 7. Three-dimensional renderings and sectional views over fractional anisotropy images show similar overall trajectories and relative pathway organization to the HCP example.

## 10. Acknowledgements

We want to thank and acknowledge Lianrui Zuo, Yihao Liu for fruitful discussions on modeling, architecture, and training.

This work was supported in part by the National Institute of Health through NIH awards K01EB032898 (KS); R01EB017230 (BL); SFP_302547 (GR). This work was supported by the Alzheimer’s Disease Sequencing Project Phenotype Harmonization Consortium (ADSP-PHC) that is funded by NIA (U24 AG074855, U01 AG068057 and R01 AG059716).

Data used in this study were obtained from the Human Connectome Project Young Adult and Human Connectome Project Development studies. The Human Connectome Project aims to map the structural connections and circuits of the brain and their relationships to behavior by acquiring high-quality magnetic resonance images.

PING: The Pediatric Imaging, Neurocognition, and Genetics (PING) dataset was collected and released openly to contribute to the assessment of typical brain development in a pediatric sample (RC2DA029475-01), (https://www.sciencedirect.com/science/article/pii/S1053811915003572).

ADNI methods: Data used in the preparation of this article were obtained from the Alzheimer’s Disease Neuroimaging Initiative (ADNI) database (adni.loni.usc.edu). The ADNI was launched in 2003 as a public-private partnership, led by Principal Investigator Michael W. Weiner, MD. The primary goal of ADNI has been to test whether serial magnetic resonance imaging (MRI), positron emission tomography (PET), other biological markers, and clinical and neuropsychological assessment can be combined to measure the progression of mild cognitive impairment (MCI) and early Alzheimer’s disease (AD).

ADNI: Data collection and sharing for the Alzheimer’s Disease Neuroimaging Initiative (ADNI) is funded by the National Institute on Aging (National Institutes of Health Grant U19 AG024904). The grantee organization is the Northern California Institute for Research and Education. In the past, ADNI has also received funding from the National Institute of Biomedical Imaging and Bioengineering, the Canadian Institutes of Health Research, and private sector contributions through the Foundation for the National Institutes of Health (FNIH) including generous contributions from the following: AbbVie, Alzheimer’s Association; Alzheimer’s Drug Discovery Foundation; Araclon Biotech; BioClinica, Inc.; Biogen; Bristol-Myers Squibb Company; CereSpir, Inc.; Cogstate; Eisai Inc.; Elan Pharmaceuticals, Inc.; Eli Lilly and Company; EuroImmun; F. Hoffmann-La Roche Ltd and its affiliated company Genentech, Inc.; Fujirebio; GE Healthcare; IXICO Ltd.; Janssen Alzheimer Immunotherapy Research & Development, LLC.; Johnson & Johnson Pharmaceutical Research &Development LLC.; Lumosity; Lundbeck; Merck & Co., Inc.; Meso Scale Diagnostics, LLC.; NeuroRx Research; Neurotrack Technologies; Novartis Pharmaceuticals Corporation; Pfizer Inc.; Piramal Imaging; Servier; Takeda Pharmaceutical Company; and Transition Therapeutics.

Data used in the preparation of this article were obtained from the Adolescent Brain Cognitive DevelopmentSM (ABCD) Study (https://abcdstudy.org), held in the NIMH Data Archive (NDA). The ABCD Study is supported by the National Institutes of Health and additional federal partners under award numbers U01DA041048, U01DA050989, U01DA051016, U01DA041022, U01DA051018, U01DA051037, U01DA050987, U01DA041174, U01DA041106, U01DA041117, U01DA041028, U01DA041134, U01DA050988, U01DA051039, U01DA041156, U01DA041025, U01DA041120, U01DA051038, U01DA041148, U01DA041093, U01DA041089, U24DA041123 and U24DA041147. A full list of supporters is available at https://abcdstudy.org/federal-partners.html. A listing of participating sites and a complete listing of the study investigators can be found at https://abcdstudy.org/consortium_members/. ABCD consortium investigators designed and implemented the study and/or provided data but did not necessarily participate in the analysis or writing of this report. This manuscript reflects the views of the authors and may not reflect the opinions or views of the NIH or ABCD consortium investigators.

BLSA: The BLSA is supported by the Intramural Research Program, National Institute on Aging, NIH. BLSA is a prospective cohort study with continuous enrollment that began in 1958. Comprehensive data from BLSA are available upon request by a proposal submission through the cohort website (www.blsa.nih.gov).

MAP/ROS/MARS: Data contributed from MAP/ROS/MARS was supported by NIA R01AG017917, P30AG10161, P30AG072975, R01AG022018, R01AG056405, UH2NS100599, UH3NS100599, R01AG064233, R01AG15819 and R01AG067482, and the Illinois Department of Public Health (Alzheimer’s Disease Research Fund). Data can be accessed at. www.radc.rush.edu. More information about participant demographics and study information can be found here: https://www.rushu.rush.edu/research-rush-university/departmental-research/rush-alzheimers-disease-center/rush-alzheimers-disease-center-research/epidemiologic-research.

The data contributed from the Wisconsin Registry for Alzheimer’s Prevention was supported by NIA AG021155, AG0271761, AG037639, and AG054047.

UpennRisk: The dataset we refer to as UPennRisk comes from a study at University of Pennsylvania that investigated whether training executive cognitive function could influence choice behavior and brain responses. Neuroimaging data were downloaded from Openneuro here: https://openneuro.org/datasets/ds002843/versions/1.0.1. We use version 1.0.1.

VKC: This work was supported by the National Institute of Child Health and Human Development (NICHD) through awards R01 HD089474 (Cutting), R37 HD095519 (Cutting), R01 HD044073 (Cutting), and R01 HD067254 (Cutting). This work was supported by the Vanderbilt Kennedy Center funded by NICHD through award P50 HD103537. This work utilized REDCap supported by the National Center for Advancing Translational Sciences (NCATS) through award UL1 TR000445 to the Vanderbilt Institute for Clinical and Translational Research. This work was supported by the National Institute of Health’s Office of the Director (1S10 OD021771-01) to the Vanderbilt University Institute of Imaging Science.

BABIES and ABC datasets were supported by the Jacobs Foundation Early Career Research Fellowship (2017-1261-05) (Humphreys); National Science Foundation CAREER Award (2042285) (Humphreys); Brain and Behavior Research Foundation John and Polly Sparks Foundation Investigator Award (29593) (Humphreys); Vanderbilt Institute for Clinical and Translational Research Grant (VR53419) (Humphreys); Vanderbilt Strong Grant; Vanderbilt Kennedy Center Grant (Humphreys); National Institute of Mental Health (R01MH129634) (Humphreys).

VUIIS-MS: This work was supported by funding from Bristol Myers Squibb and NIH 5R01NS117816-04 and K01EB032898.

HBN: The Healthy Brain Network (HBN) is an ongoing initiative focused on building a biobank of data from 10,000 children and adolescents (ages 5-21) in the New York City area (https://www.nature.com/articles/sdata2017181). Phenotypic data are available upon request by filling out a data use agreement: https://fcon_1000.projects.nitrc.org/indi/cmi_healthy_brain_network/Phenotypic.html.

VMAP/TAP (and AJ): Alzheimer’s Association IIRG-08-88733 (ALJ); R01-AG034962 (ALJ); R01-AG056534 (ALJ); K24-AG046373 (ALJ); UL1-TR000445 and UL1-TR002243 (Vanderbilt Clinical Translational Science Award); P20-AG068082 and P30-AG086403 (Vanderbilt Alzheimer’s Disease Research Center); S10-OD023680 (Vanderbilt’s High-Performance Computer Cluster for Biomedical Research) ; S10-OD021771 (VUIIS Center for Human Imaging); Richard Eugene Hickman Alzheimer’s Disease Research Endowment; Vanderbilt Memory & Alzheimer’s Center

